# Increased contact transmission of contemporary Human H5N1 compared to Bovine and Mountain Lion H5N1 in a hamster model

**DOI:** 10.1101/2025.06.23.661180

**Authors:** Reshma Koolaparambil Mukesh, Franziska Kaiser, Jonathan E. Schulz, Shane Gallogly, Jessica Prado-Smith, Arthur Wickenhagen, Kathleen Cordova, Brian J Smith, Chad Clancy, Carl Shaia, Greg Saturday, Emmie de Wit, Neeltje van Doremalen, Claude Kwe Yinda, Vincent J Munster

**Affiliations:** Laboratory of Virology, National Institute of Allergy and Infectious Diseases, National Institutes of Health, Hamilton, MT, USA; Rocky Mountain Veterinary Branch, National Institute of Allergy and Infectious Diseases, National Institutes of Health, Hamilton, MT, USA

## Abstract

The ongoing outbreak of highly pathogenic avian influenza virus (HPAIV) subtype H5N1 in the U.S. poses a significant public health threat. To date, 70 human cases have been confirmed in the United States, including two severe cases and one fatality. While suitable animal models are crucial for predicting the potential pandemic risk of newly emerging pathogens in humans, studies investigating contemporary HPAIV H5N1 transmission dynamics remain limited. Here, we investigated the pathogenicity and transmission efficiency of three recent clade 2.3.4.4b H5N1 viruses isolated from a bovine, mountain lion, and a human case using Syrian hamsters. Intranasal inoculation with 10^4^ TCID_50_ resulted in productive virus replication in the respiratory tract and shedding for all three isolates. Transmission studies showed limited efficiency via direct contact and airborne routes for all three isolates. Although overall transmission was inefficient, the human H5N1 isolate demonstrated relatively greater contact transmissibility than the bovine and mountain lion isolates. Taken together, our data demonstrate that the Syrian hamster model complements existing animal models for influenza A virus research and expands the resources available for investigating the pathogenicity, transmissibility, and efficacy of countermeasures against HPAIV H5N1.

**One sentence summary:** Human HPAIV H5N1 exhibits comparatively higher contact transmissibility than bovine and mountain lion isolates, despite limited overall transmission.

## Introduction

Over the last decades, Highly Pathogenic Avian Influenza virus (HPAIV) H5N1 has evolved from a virus circulating mainly in poultry in Southeast Asia into a panzootic virus with circulation on all continents, including Antarctica (*1–4*). The introduction of the clade 2.3.4.4b HPAIV H5N1 into the U.S. in 2021 was followed by rapid dissemination via migratory birds and spillover into a wide variety of mammals, backyard poultry and commercial poultry operations (*4–6*).

In the United States, until recently, the main mammalian species in which HPAIV H5N1 was detected were domestic and wild carnivorous species, including domestic cats (*Felis catus*), black bears (*Ursus americanus*), bobcats (*Lynx rufus*), red foxes *(Vulpes vulpes*), harbor seals *(Phoca vitulina*), and mountain lions (*Puma concolor*) (*7*). In February 2024, the first detection of HPAIV H5N1 clade 2.3.4.4b, genotype B3.13 in dairy cattle was observed in Texas; from this initial introduction, the virus has spread to 17 states and has affected over 1073 dairy herds (as of June 12, 2025) (*8*). Within cattle, HPAIV H5N1 appears to be mainly replicating in the mammary tissue (*9*). Transmission between dairy cattle is thought to occur via a combination of routes, including fomite transmission via contaminated milking equipment (*10*). HPAIV H5N1 has been detected by PCR in milk within the food chain; however, several studies have confirmed the rapid inactivation of the virus by commonly used pasteurization methods (*11–13*). In January 2025, a second independent introduction of HPAIV H5N1, clade 2.3.4.4b, genotype D1.1, in dairy cattle was identified in Nevada (*14*).

The significant increase of HPAIV H5N1 in the animals and environment resulted in spillover into humans. To date, 70 human cases have been reported, with most traced back to exposure to either dairy cattle or poultry (*15*). In a few instances, the exposure history was unknown. HPAIV H5N1 in humans in the U.S. has generally caused mild illness, including conjunctivitis (*16*). However, in 2025, the first fatal HPAIV H5N1 infection in the U.S. was reported, caused by a H5N1 clade 2.3.4.4b, genotype D1.1 (*17*). In several human cases, the HPAIV H5N1 exhibited markers of human adaptation including the E627K in PB2, and E186D and Q222H in the HA gene segments (*18*).

Currently, the clade 2.3.4.4b HPAI H5N1 viruses circulating in cows have retained their primary avian-type receptor preference (*19*), potentially due to the abundant expression of α2,3-linked sialic acid in the mammary glands of dairy cattle (*20*). A limited number of amino acid substitutions in the H5 HA are believed to shift receptor binding preference from avian-type to human-type α2,6-linked sialic acids, a change associated with increased airborne transmission (*21–23*). Despite this avian-type receptor preference of the HPAI H5N1 clade 2.3.4.4b viruses, efficient contact and airborne transmission have been observed in the ferret model, a model typically used to assess the human-to-human transmission efficiency of influenza A viruses (*24–26*). To date, there is no evidence of human-to-human transmission associated with the ongoing outbreak in the United States.

Syrian hamsters have historically been developed as a small animal model for influenza A virus, and more recently as a model to study transmission (*27–31*). H1N1 and H3N2 seasonal influenza A viruses are transmitted via the airborne route, as observed in the ferret model (*32–34*). During the COVID-19 pandemic, the Syrian hamster model played a crucial role in understanding the drivers of human-to-human transmission of SARS-CoV-2 (*35, 36*). In our study, we demonstrated the use of the Syrian hamster model to evaluate the transmission potential of three contemporary HPAIV H5N1 isolates obtained from a bovine, mountain lion and a human case: A/bovine/Ohio/B24OSU-342/2024, clade 2.3.4.4b, genotype B3.13 (Designated as Bovine), A/mountain lion/Montana/1/2024, clade 2.3.4.4b, genotype B3.6 (Designated as Mountain Lion) obtained from the lung tissue of a deceased mountain lion, and A/Texas/37/2024, clade 2.3.4.4b, genotype B3.13 (Designated as Human). We inoculated hamsters intranasally (I.N.) with 10^4^ TCID_50_ of each of the three HPAIV H5N1 isolates and monitored disease progression and survival. Shedding kinetics and transmission efficiency were assessed for the direct contact and airborne routes using virus detection and seroconversion. The Human isolate demonstrated productive transmission, with two of eight direct contact sentinels exhibiting active virus replication and shedding. In contrast, the Bovine and Mountain Lion isolates resulted in unproductive direct contact transmission in one of eight sentinels, with seroconversion but no virus detection. Unproductive airborne transmission was also observed in one of eight sentinels exposed to the Bovine isolate, resulting in seroconversion without virus detection. Our data demonstrate that, despite high levels of virus replication in the upper respiratory tract of hamsters, the efficiency of the Bovine, Mountain Lion and Human HPAIV H5N1 isolates to transmit via direct contact and airborne routes is limited.

## Results

### Productive infection of Syrian hamsters with contemporary HPAIV H5N1 isolates

To better understand the amino acid level differences among the three HPAIV H5N1 isolates (Bovine, Mountain Lion, and Human), we aligned the sequence of the respective gene segments. A summary of the amino acid variation among the three isolates is provided in Table S1. Among the three isolates, the PB2 E627K substitution was found exclusively in the Human isolate.

To evaluate the fitness of three contemporary HPAI H5N1 viruses *in vivo*, we inoculated Syrian hamsters (n=12) I.N. with 10^4^ TCID_50_ of each isolate and assessed replication and shedding kinetics (Fig. 1A). Six animals per group were scheduled for necropsy at four days post inoculation (dpi) to determine virus titers in the respiratory tissue samples. The remaining six animals per group were monitored daily for disease progression and survival.

**Figure 1.**
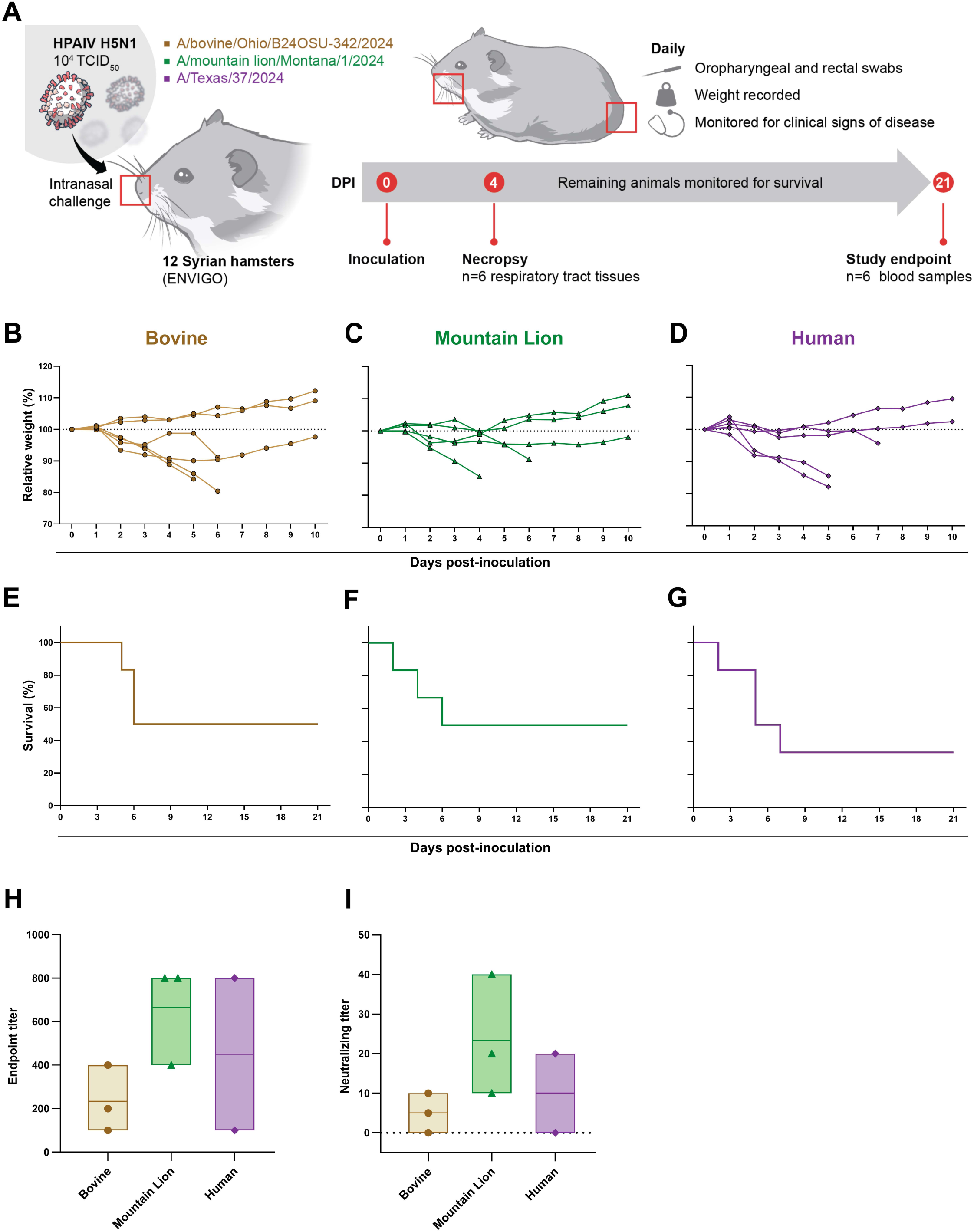
Productive infection of Syrian hamsters with contemporary H5N1 influenza A viruses. **(A)** Schematic overview of the experimental design. Syrian hamsters (n=12 per group) were I.N. inoculated with 10□ TCID□□ of either A/bovine/Ohio/B24OSU-342/2024 (Bovine), A/mountain lion/Montana/1/2024 (Mountain Lion), or A/Texas/37/2024 (Human) in 40 µL of sterile DMEM. At 4 dpi, six animals from each group were euthanized for collection of respiratory tissues. The remaining six animals per group were monitored daily for clinical signs, weight loss, and survival up to 21 dpi. Oropharyngeal and rectal swabs were collected daily for 10 days. **(B-D)** Relative body weight changes over a 10-day period in hamsters infected with Bovine (B), Mountain Lion (C), or Human (D) HPAIV H5N1 isolates (n=6 per group). **(E–G)** Kaplan–Meier survival curves for each group. Survival proportions were compared using the Log-rank (Mantel-Cox) test. **(H, I)** Endpoint titers of humoral IgG anti-H5 HA responses in surviving animals at 21 dpi by ELISA (H), and corresponding virus-neutralizing titers measured by microneutralization assay (I). Individual values are shown, bars represent the range (minimum to maximum), with the middle line indicating the median.

Clinical signs of disease appeared from 2 dpi onward and included ruffled fur, hunched posture, hypoactivity, open-mouth breathing, dyspnea, and respiratory distress in all three groups. In the Bovine group, four animals began losing weight from 2 dpi. Of these, three reached predetermined euthanasia endpoint criteria on days five and six, while the fourth started to regain weight from day six onward (Fig. 1B). One animal from each of the Mountain Lion and Human groups succumbed to disease on day 2. Additionally, two animals from the Mountain Lion group and three from the Human group met euthanasia criteria between days four and seven (Fig. 1C & D). Overall, the survival after 21 dpi was 50% (3/6) in the Bovine (Fig. 1E) and Mountain Lion (Fig. 1F) groups and 33.33% (2/6) in the Human group (Fig. 1G). All surviving animals were seroconverted; however, endpoint antibody titers varied. The Bovine group had endpoint titers between 100 and 400, whereas the Mountain Lion group had titers ranging from 400 to 800. The two animals in the Human group had endpoint titers of 100 and 800 (Fig. 1H). Consistent with these results, neutralizing antibody titers were highest in the Mountain Lion group, whereas the Bovine and Human groups exhibited comparatively lower neutralizing titers (Fig. 1I).

### Hamsters inoculated with Human HPAIV H5N1 shed more infectious virus compared to Bovine and Mountain Lion H5N1 isolates

Oropharyngeal and rectal swabs were collected daily to determine the kinetics of virus shedding. High viral RNA loads were detected by qRT-PCR in the oropharyngeal swab samples of all animals from all three groups with relatively similar shedding kinetics, viral RNA levels started to decline from 5 dpi onwards and no viral RNA was detected after 8 dpi (Fig. 2A). No significant differences were observed in total amounts of viral RNA shed as analyzed by area under the curve analysis (Fig. 2B). Corresponding to the amount of virus RNA, infectious virus was also found to be high during the earlier days of infection specifically on 1 and 2 dpi and started to decline thereafter (Fig. 2C). Significant differences were observed in the total amount of infectious virus shed, with the Human group shedding more infectious virus compared to the Mountain Lion group (Fig. 2D). Although viral RNA was detectable in rectal swabs, it was not consistently detected in all animals within each group (Fig. 2E). Analysis of the area under the curve revealed no significant differences in the total viral RNA shed from the intestinal tract among the three groups (Fig. 2F). In addition, infectious virus could only be found in one animal from the Bovine group (Fig. 2G), indicating that shedding from the intestinal tract is not prominent compared to shedding from the respiratory tract.

**Figure 2.**
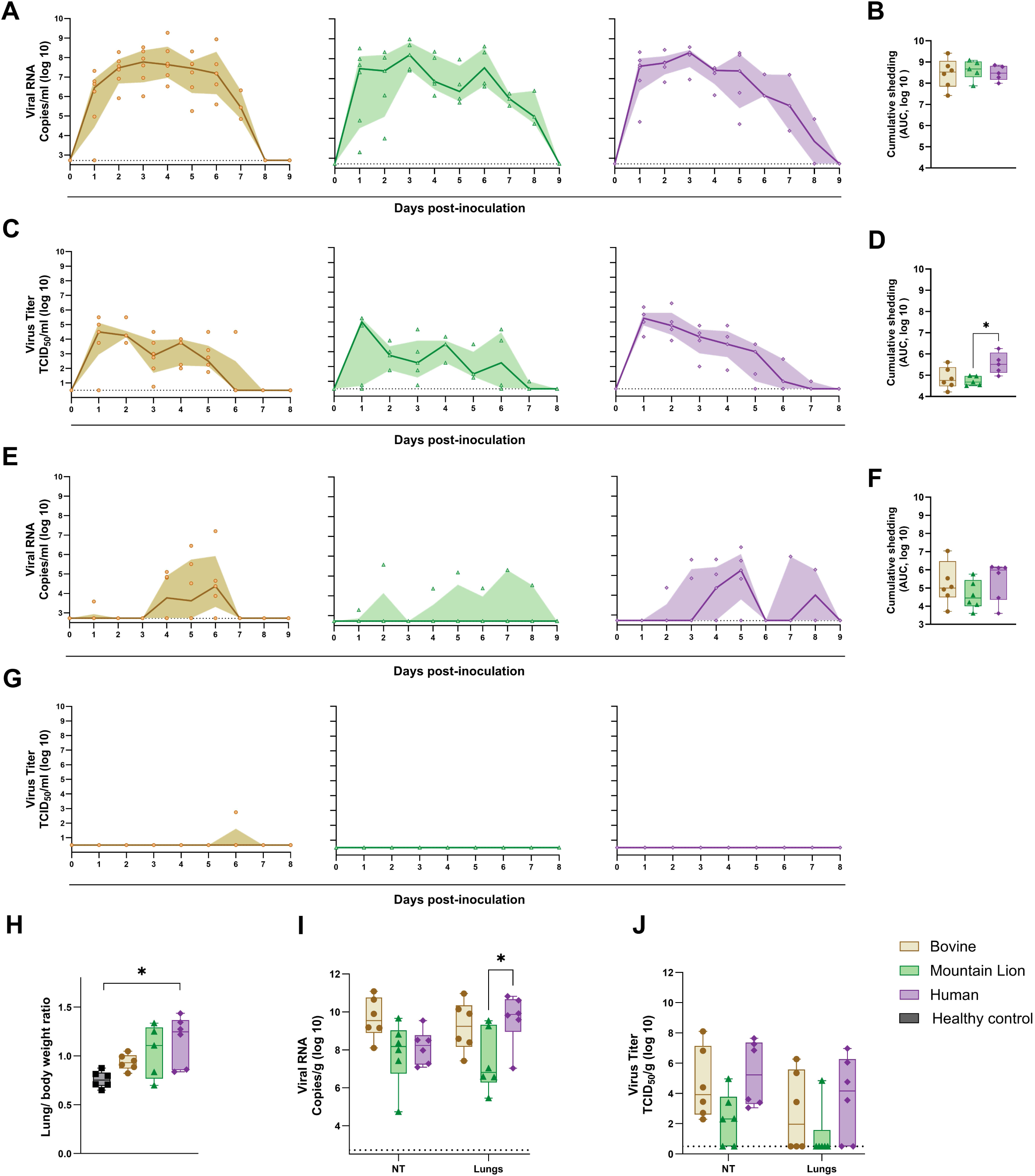
Hamsters inoculated with Human HPAIV H5N1 isolate shed more infectious virus compared to Bovine and Mountain Lion H5N1 isolates. (A, **B)** Viral shedding kinetics as measured as RNA copy number in oropharyngeal swabs and corresponding area under the curve (AUC) analyses. Oropharyngeal swabs were collected daily and viral RNA was quantified by RT-qPCR. The AUC of viral RNA shedding was calculated for each group. **(C, D)** Quantification of infectious HPAIV H5N1 in oropharyngeal swabs and corresponding AUC analysis. Infectious virus was measured by endpoint titration, and AUC was calculated to compare total oral shedding. **(E, F)** Viral RNA levels in rectal swabs and corresponding AUC analysis, as measured by RT-qPCR. **(G)** Infectious virus titers in rectal swabs. **(H)** Lung-to-body weight ratios at 4 dpi. Lungs were harvested from infected and uninfected control animals (n=6 per group), and lung weight to body weight ratio was determined to assess pulmonary pathology. **(I, J)** Nasal turbinates (NT) and lung tissues collected at 4 dpi were analyzed for viral RNA (I) and infectious virus (J). Statistical significance was assessed using a Kruskal-Wallis test followed by the Mann–Whitney U test for B, D and F; Kruskal-Wallis test followed by Dunn’s multiple comparisons test for H; and two-way ANOVA followed by Tukey’s multiple comparisons test for I and J. * = p-value <0.05. Limit of detection was 2.72□log_10_ Copies/mL for A, E, and I, 0.5□log_10_ TCID_50_/mL for C and G and 0.5 log_10_ TCID_50_/g for J. For panels A, C, E, and G, individual values are shown; the line represents the median, and the shaded area indicates the interquartile range. For panels B, D, F, and H-J, individual values are shown; bars represent the range (minimum to maximum), and the middle line indicates the median (n=6).

To determine virus replication in the upper and lower respiratory tract tissues, nasal turbinates (NT) and lungs were collected at 4 dpi. For six animals of each group, the relative lung/body weight was measured at necropsy on 4 dpi as an indicator of lung inflammation. In comparison with healthy control animals, a significant increase in relative lung weight was observed in the Human group (Fig. 2H). Virus replication was detected in both the upper and lower respiratory tracts of all animals across the three groups. Although not significant, animals in the Bovine group had a relatively higher amount of viral RNA in the NT in comparison with Mountain Lion and Human groups (Fig. 2I). The amount of viral RNA in the lungs of infected animals was significantly higher in the Human group compared to the Mountain Lion group.

A similar pattern was observed for infectious virus levels in the lungs, with the Bovine and Human groups exhibiting, on average, higher viral loads compared to the Mountain Lion group (Fig. 2J). However, no significant difference in the amount of infectious virus in the NT and lungs was found between the three groups. Whereas high amounts of viral RNA were detected in the lung tissues of infected animals from all three groups, infectious virus could only be recovered from a subset of these animals in the Bovine (3/6), Mountain Lion (1/6) and Human (4/6) groups. Overall, the Bovine and Human H5N1 viruses replicated to higher titers in the respiratory tract of the hamsters compared to the Mountain Lion H5N1 virus.

### More pronounced histopathologic lesions in the respiratory tract with Bovine and Human H5N1s compared to Mountain Lion isolate

While all three HPAIV H5N1 isolates replicated efficiently in the upper and lower respiratory tract of the hamsters, we were interested in whether the different viruses would display a differential disease phenotype and lung pathology. To better understand potential differences in disease phenotype, four animals per group were inoculated with 10□ TCID□□ of each virus and euthanized at 4 dpi. Respiratory tract tissues including NT, trachea, and lungs were collected for comprehensive histopathological evaluation of the entire respiratory tract. Histopathological evaluation revealed findings consistent with viral infection in hamsters (Table S2). The NT showed no inflammatory response in any of the groups, and minimal necrosis was observed in only two out of four animals from the Human group (Fig. 3A & B). Tracheal samples from all three groups contained varying amounts of degenerate or necrotic epithelial cells, along with abundant luminal exudate composed of sloughed necrotic debris. Slightly more pronounced necrosis and neutrophilic tracheitis were observed in the Bovine and Human groups (Fig. 3A & C). Tracheal lesions were more pronounced in the Bovine (n=4/4) and Human groups (n=4/4). In contrast, the Mountain Lion group (n=2/4) had only two animals with minimal to moderate tracheal lesions (Fig. 3C). Pulmonary lesions were characterized by necrotizing broncho-interstitial pneumonia, consisting of multifocal inflammatory nodules centered on terminal bronchioles and extending into adjacent alveoli (Fig. 3A). The bronchiolar epithelium was frequently degenerate or necrotic, with abundant luminal exudate. These inflammatory nodules were composed predominantly of foamy macrophages, with fewer neutrophils and lymphocytes, and small amounts of necrotic debris. In most cases, hemorrhage, fibrin, and edema were present and intermingled with inflammatory cells, often extending into the surrounding alveoli. Adjacent alveoli were thickened due to the presence of fibrin, edema, and a small number of macrophages and neutrophils. Pulmonary pathology was more pronounced in the Bovine and Human groups. The Bovine group (n=3/4) had a mild to marked distribution of lesions while the Human group (n=2/4) had mild lesions (Fig. 3D). Pulmonary lesions were not observed in the Mountain Lion group. Overall, the Mountain Lion group exhibited comparatively mild histopathological lesions in the respiratory tract relative to the Bovine and Human groups.

**Figure 3.**
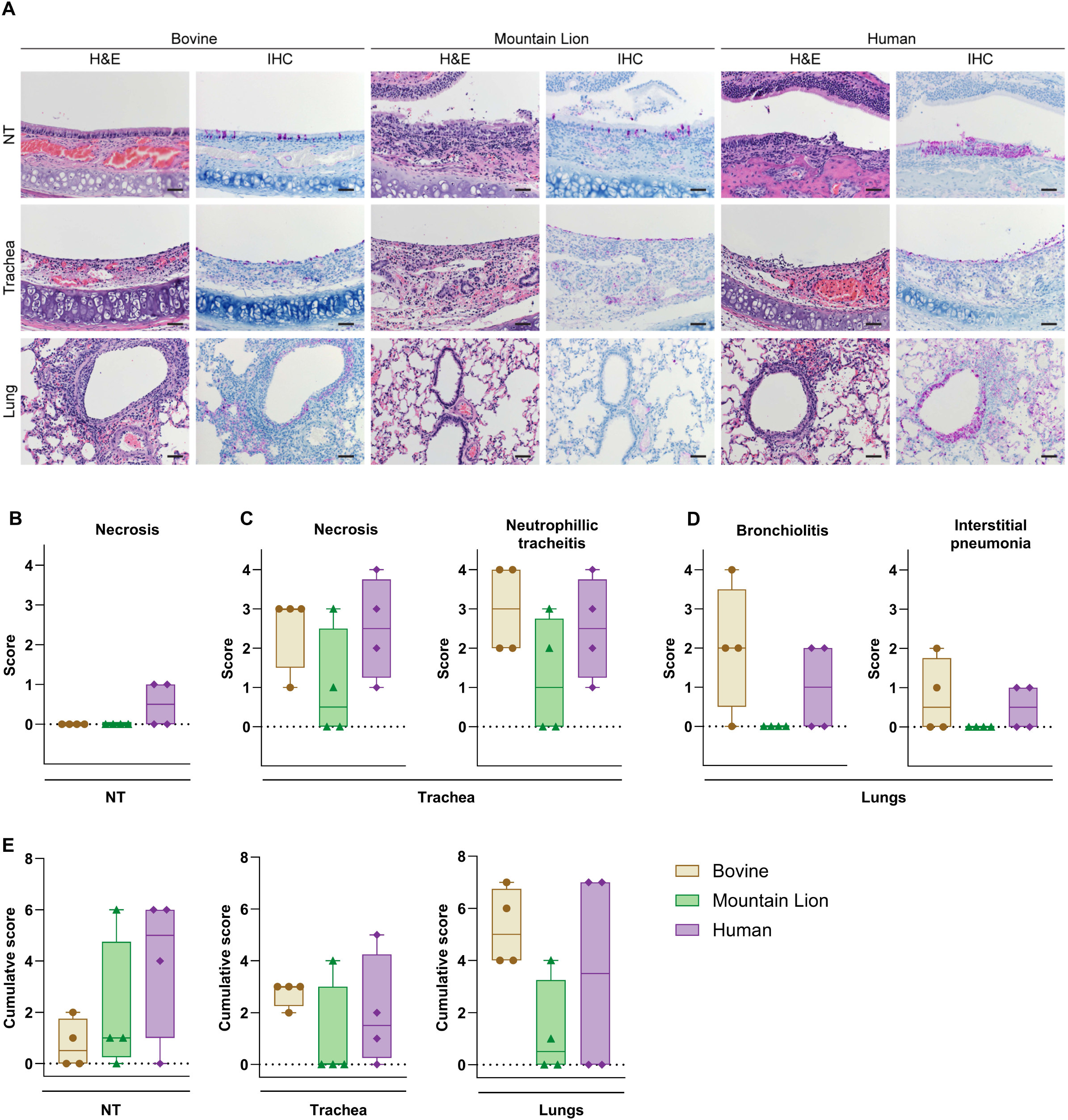
More pronounced histopathologic lesions in the respiratory tract with Bovine and Human H5N1s compared to Mountain Lion isolate. **(A)** Representative images of hematoxylin and eosin-stained (H&E) and immunohistochemistry (IHC) of NT, trachea, and lung tissues collected at 4 dpi. Influenza A virus antigen was detected by IHC targeting influenza A virus nucleoprotein. Magnification and scale bars: Nasal turbinates: 400×, bar = 20 µm, Trachea: 200×, bar = 50 µm, and Lungs: 200×, bar = 50 µm. **(B-D)** Semi quantitative pathological scoring of NT (B), trachea (C), and lungs (D). **(E)** Cumulative scoring of the presence of viral antigen in the respiratory tissues based on IHC staining. Statistical significance for panels B-E was assessed using the Kruskal-Wallis test. For graphs B-E, individual values are shown; bars represent the range (minimum to maximum), with the middle line indicating the median (n=4).

To evaluate cellular tropism, immunohistochemistry (IHC) was performed to detect Influenza A nucleoprotein as a marker of virus replication. Viral antigen was present mostly on the respiratory and olfactory epithelial cells of the upper respiratory tract (Table S2). Cumulative IHC on the NT showed more staining in the Human group (3/4) compared to the Bovine and Mountain Lion groups (Fig. 3A & E). In the trachea, replication was mostly observed in the ciliated epithelial cells and cumulative IHC showed more staining in the Bovine and Human groups compared to the Mountain Lion group (Fig. 3A & E). Immunohistochemical evaluation of the lower respiratory tree revealed the presence of viral antigen in the bronchiolar epithelium, pneumocytes and alveolar macrophages (Table S2). Viral antigen could be detected in 4/4 hamsters in the Bovine, 2/4 hamsters in the Mountain Lion and 2/4 hamsters in the Human groups (Fig. 3A & E). Although the pathogenic phenotypes were overall comparable between the groups, slightly more influenza A virus replication was observed by IHC within the respiratory tract of hamsters in the Bovine and Human groups compared to the Mountain Lion group.

### Productive transmission of the Human HPAIV H5N1 via direct contact but not the airborne route

Recent analyses of the transmission kinetics of contemporary HPAI H5 viruses in ferrets have demonstrated increased viral shedding into the air, with efficient direct contact and partial airborne transmission (*24, 37*). However, within the current outbreak of Bovine HPAIV H5N1 in the U.S., no evidence of human-to-human transmission has been observed, and reported human cases have primarily resulted from exposure to infected livestock or poultry (*38*). In this study, we aimed to investigate the transmission potential of three different HPAIV H5N1 isolates using the hamster model to complement existing data generated in the ferret model.

Eight donor animals per group were inoculated I.N. with 10^4^ TCID_50_ of each virus. The transmission cage set-up used a cage divider, which allowed us to study direct-contact and airborne transmission within the same cage (*36*). On day 0, inoculated donor animals were placed in the cage and 24 hours thereafter naive direct-contact and airborne sentinel hamsters were introduced into the transmission cage. A total of eight individual transmission pairs were used to test the transmission efficiency. At the end of the 48-h transmission window (days 1 to 3 post-inoculation of the donors), the contact and airborne sentinel hamsters were single-housed, oropharyngeal swabs were collected daily and monitored for signs of disease. To assess transmission efficiency, detection of virus replication (viral RNA from the oropharyngeal swab samples) and seroconversion on day 21 after exposure (the presence of binding antibodies against H5) were used. All donor animals from the three groups were productively infected and shed viral RNA during the transmission window (Fig. 4A-C).

**Figure 4.**
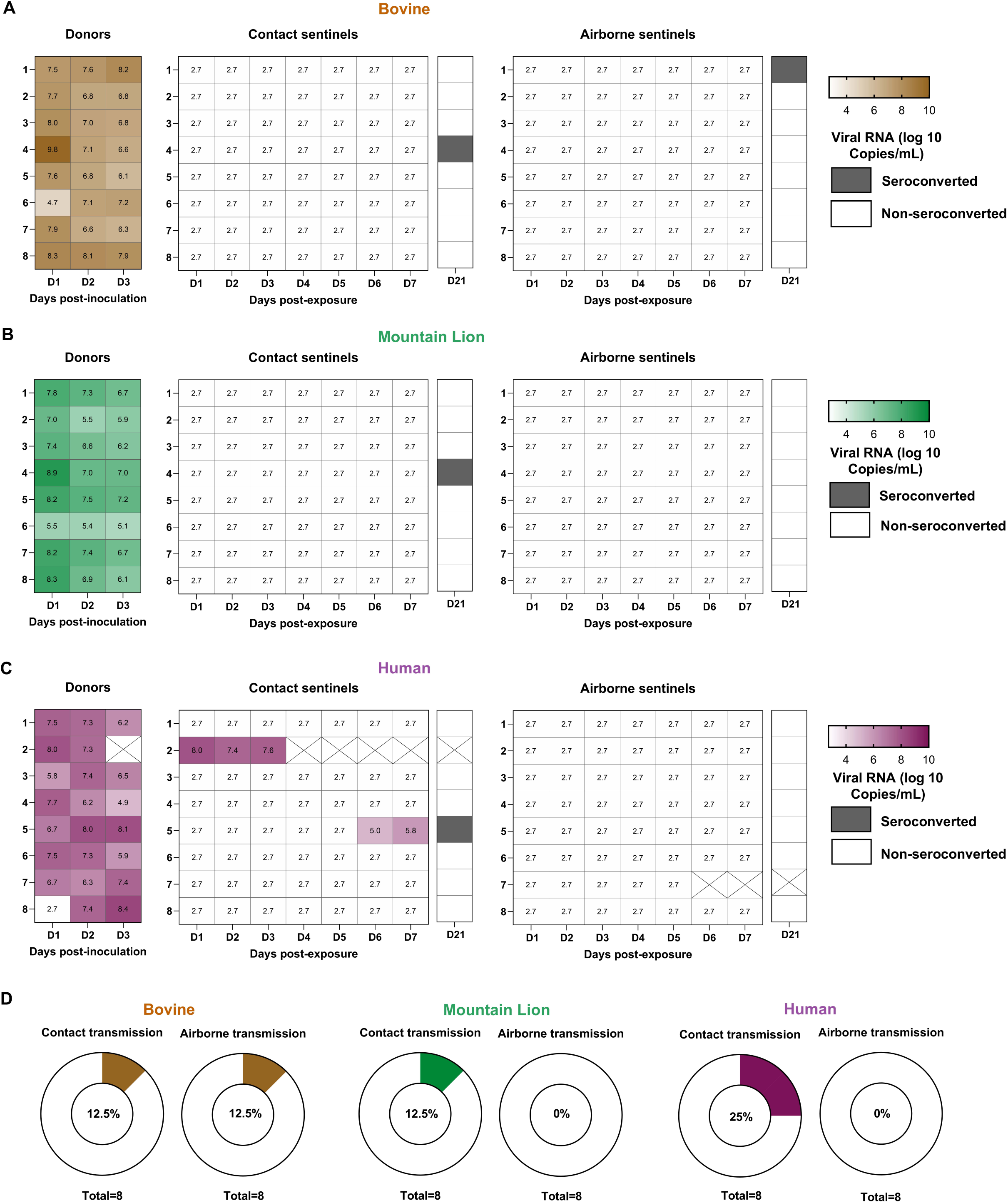
Productive direct contact, but not airborne transmission of Human HPAIV H5N1 (A-C) Shedding kinetics of viral RNA and seroconversion data obtained after transmission study. Donor hamsters (n=8 per group) were intranasally inoculated with 10□ TCID□□ of one of three HPAIV H5N1 isolates: Bovine, Mountain Lion, or Human. At 1 dpi of donor animals, naïve sentinel animals were introduced into the transmission cages to assess virus transmission via either direct contact (n=8 per transmission group) or airborne exposure (n=8 per transmission group). Transmission was evaluated based on detection of viral RNA in oropharyngeal swabs and/or seroconversion at 21 dpi. (C) Summary of overall transmission efficiency. Transmission efficiency via direct contact and airborne routes for each virus is presented as donut plots. Limit of detection was 2.72□log_10_ Copies/mL for A-C.

Despite the high levels of viral shedding from donor animals in all three groups, productive transmission via the direct contact route was observed only in the Human group (Fig. 4C). Two out of eight animals became productively infected, with high viral loads, similar to the donor animals, shed from the respiratory tract. Contact sentinel animal 2 in the Human group began showing signs of disease on 2 days post-exposure and reached euthanasia endpoint criteria on day 4. The remaining infected contact sentinel animal (Sentinel 5) in the Human group showed no severe signs of disease despite viral shedding and seroconverted.

Contact sentinel 4 in both the Bovine and Mountain Lion groups seroconverted without showing any clinical signs of infection and no detection of viral RNA in oropharyngeal swabs. This suggests that virus transmission occurred but did not result in a productive infection, as it was quenched rapidly after transmission. Overall, the transmission efficiency via direct contact was 12.5% for the Bovine and Mountain Lion groups, and 25% for the Human isolate group (Fig. 4D).

Transmission via the airborne route was detected exclusively in the Bovine group (Fig. 4A). Sentinel animal 1 from the Bovine group seroconverted without detection of viral RNA in the oropharyngeal swabs, suggests that virus transmission did not result in a productive infection, but was quenched rapidly after transmission, comparable to the contact transmission of the Bovine group. Taken together, our results demonstrate that the Human HPAIV H5N1 has a higher transmission potential compared to the Bovine and Mountain Lion isolates, as productive transmission was observed only with the Human isolate.

## Discussion

The zoonotic and human-to-human transmission potential of HPAIV H5Nx viruses has been under investigation since the first human cases were reported in 1997 in Hong Kong (*39*). The introduction of HPAIV H5N1 in the Americas and subsequent outbreaks in poultry and dairy cattle have dramatically increased human exposure to these viruses (*40, 41*). The majority of human cases in the U.S. stem either from exposure through infected poultry or dairy cattle. However, sustained human-to-human transmission of HPAIV H5N1 has so far not been reported (*42*).

Since the first detection of the HPAIV H5N1 clade 2.3.4.4b in dairy cattle, several studies have investigated the pathogenicity and transmission potential of HPAIV H5N1 isolates using a variety of animal models including mice, ferrets and non-human primates. Among the variety of mouse models utilized, the HPAIV H5N1 typically causes rapid generalized disease with systemic replication and uniformly lethal outcomes, regardless of the mouse strain or exposure route (*24, 43, 44*). In addition, 10^6^ plaque-forming units (PFU) of the Bovine and Human H5N1 isolates caused severe disease in ferrets resulting in rapid disease progression, systemic replication, and uniform lethality (*24, 25*). The pathogenicity associated with 10^7^ TCID_50_ of the Bovine H5N1 isolate in non-human primates was dependent on the inoculation route. While intratracheal inoculation resulted in severe, fatal broncho-interstitial pneumonia, intranasal or orogastric exposure led to mild or subclinical disease. Regardless of the route of inoculation, all animals shed virus both orally and nasally (*45*).

While hamsters are a well-established small animal model for influenza A virus research, they are not as widely used as ferrets or Guinea pigs (*27, 46*). Hamsters have been extensively used to evaluate the transmission of several respiratory pathogens, including SARS-CoV-2 and Nipah virus, making them an attractive animal model for investigating transmission (*35, 36, 47*). A recent hamster study evaluated the pathogenicity and transmission potential of the Human H5N1 isolate using 10^3^ PFU in 30 µl of PBS, and demonstrated that hamsters are susceptible to HPAIV H5N1, showing high lethality and 100% transmission efficiency via direct contact (*48*).

In our study, intranasal inoculation with 10^4^ TCID□□ resulted in 50% survival in both the Bovine and Mountain Lion groups and 33.3% survival in the Human group. Infected hamsters displayed differences in shedding and replication between the three isolates. Interestingly, cumulative respiratory shedding of viral RNA did not differ significantly among the three groups. However, higher levels of infectious virus were detected in the Human group compared to the Mountain Lion group, suggesting potential differences in viral replication efficiency or host-pathogen interactions despite similar RNA shedding profiles. These findings are consistent with the observed trends in viral RNA levels in the URT, where the Bovine group exhibited higher RNA loads, while comparable levels were detected between the Human and Mountain Lion groups. Notably, despite similar RNA loads, differences in infectious virus titers in the oropharyngeal swab samples were observed only between the Human and Mountain Lion groups, indicating potential differences in viral replication competence or shedding of infectious particles. Although these viruses replicated efficiently and were shed from the respiratory tract of hamsters, their relative transmission efficiency remained low. This discrepancy may be attributed to the inefficient release of infectious viral particles into the air, limiting onward transmission. Among the three isolates tested, productive transmission was observed only with the Human isolate. This observation is supported by a recent study in ferrets which displayed a higher release of infectious virus in the air with the Human HPAIV H5N1 than Bovine HPAIV H5N1 (*25, 49*).

The Human isolate, A/Texas/37/2024, has now been utilized in a wide variety of *in vitro* and *in vivo* studies, resulting in a better understanding of the phenotypic characteristics of this virus. The A/Texas/37/2024 isolate contains the PB2-E627K substitution, a mutation known to enhance the replication capacity of influenza viruses in mammalian hosts (*50*). In A/Texas/37/2024, the PB2-E627K increases the viral polymerase activity in minigenome assays (*37*). The PB2-E627K substitution is a common occurrence and has been identified in multiple human HPAIV H5N1 cases, independent of exposure source (e.g. dairy cattle or poultry), or lineage (Clade 1, Clade 2.2, and Clade 2.3.4.4b) (*51, 52*). In our study, the Bovine and Human isolates exhibited a more pathogenic phenotype compared to the Mountain Lion isolate. Although all three viruses replicated throughout the upper and lower respiratory tract, the Bovine and Human isolates induced more extensive pulmonary lesions compared to the Mountain Lion isolate. This difference may be attributable to the distinct genetic makeup of the Mountain Lion isolate (genotype B3.6).

Historic and contemporary HPAIV H5N1 isolates have been extensively characterized in the ferret model for their potential to transmit through direct contact and airborne routes. In the ferret model, the Bovine isolate did not transmit efficiently via the airborne route, whereas the Human isolate transmitted efficiently via the direct contact and airborne transmission routes (*24, 25*). Typically, transmission events resulted in fatal disease in recipient ferrets. Within our study, only direct contact transmission with the Human isolate resulted in productive infection and seroconversion. In contrast, one direct contact sentinel each from the Bovine and Mountain Lion groups seroconverted without detectable replicating virus in oropharyngeal swabs. For the airborne route, only a single sentinel in the Bovine group was seroconverted, again without the detection of replicating virus in the oropharyngeal swabs. Our results are in agreement with a recent study that only used the Human isolate and displayed contact but not airborne transmission in the hamsters, albeit at a higher efficiency than in our study (*48*).

The relatively inefficient transmission in our study aligns with the lack of reported human-to-human transmission. Comparable to other studies, increased viral shedding with the Human isolate is associated with a higher transmission potential. Despite the productive infection in the upper respiratory tract of infected hamsters, the relatively low transmission efficiency suggests that additional phenotypic changes are necessary for increased transmission. The relative receptor distribution pattern in hamsters is comparable to that in other species, including humans, with the expression of both α2,6- and α2,3-linked sialic acid receptors in the respiratory tract, albeit at different ratios (*27*). Recent studies have demonstrated that the Bovine and Human isolates maintain avian-type receptor-binding specificity (*53*). In addition, *in vitro* studies have shown that the Q226L and N224K substitutions in the HA of previous human HPAIV H5N1 isolates shift receptor specificity toward α2,6-linked sialic acids, potentially enhancing the transmission potential. This observation aligns with findings from the ferret model, where alterations in receptor specificity have been linked to efficient airborne transmission of HPAIV H5N1 (*21–23*).

The ongoing circulation of HPAIV H5 in wild birds, wild mammals, poultry and dairy cattle is cause for considerable concern. The continued circulation of these viruses significantly heightens the risk of adaptations that could enhance zoonotic potential and facilitate more efficient human-to-human transmission. Similar to the SAVE program established for SARS-CoV-2, real-time risk assessment of emerging mutations in HPAIV H5N1 is essential to evaluate their potential impact on pathogenicity, transmissibility, and the effectiveness of available countermeasures (*54*). The hamster model complements established animal models, such as the ferret model, and broadens the available tools for studying the pathogenicity and transmission dynamics associated with HPAIV H5N1 infection.

## Materials and Methods

### Ethics statement

All animal experiments were conducted after obtaining prior approval from the Institutional Animal Care and Use Committee of Rocky Mountain Laboratories, National Institutes of Health. Experiments were carried out in an Association for Assessment and Accreditation of Laboratory Animal Care International-accredited facility, following the guidelines and basic principles in the Guide for the Care and Use of Laboratory Animals, the Animal Welfare Act, US Department of Agriculture, and the US Public Health Service Policy on Humane Care and Use of Laboratory Animals. Syrian hamsters of age >4 weeks were used in this study. All work with infectious HPAI H5N1 viruses was approved for biosafety level 3 (BSL-3) conditions by the Institutional Biosafety Committee (IBC). All sample inactivation procedures were carried out in accordance with IBC-approved standard operating procedures for the removal of specimens from high-containment facilities.

### Virus and cells

Human HPAIV H5N1 isolate A/Texas/37/2024 (EPI_ISL_19027114) was obtained from Dr. Todd Davis at the Centers for Disease Control and Prevention, Decatur, Georgia, USA. Bovine HPAIV H5N1 isolate A/bovine/Ohio/B24OSU-342/2024 (EPI_ISL_19178076) was obtained from Richard Webby at St. Jude’s Children hospital, Memphis, TN, USA and Andrew Bowman at Ohio State University, Columbus, OH, USA. Mountain lion HPAIV H5N1 isolate A/mountain lion/Montana/1/2024 (EPI_ISL_19083124) was obtained from a lung sample of a diseased mountain lion in Montana in February 2024. All three viruses belong to the 2.3.4.4b clade of HPAIV H5N1. The Bovine and Human isolates are classified as genotype B3.13, whereas the Mountain Lion isolate belongs to genotype B3.6.

Virus propagation was performed in Madin-Darby canine kidney (MDCK) cells in MEM supplemented with 1□mM L-glutamine, 50□U/mL penicillin, 50□μg/mL streptomycin, 20 mM HEPES, and 4 µg/mL TPCK trypsin. MDCK cells were maintained in MEM supplemented with 10% fetal bovine serum, 1□mM L-glutamine, 50□U/ml penicillin, and 50□μg/ml streptomycin, 20 mM HEPES. Mycoplasma testing is performed at regular intervals. No mycoplasma was detected during the study.

### Animal studies

Twelve male and female Syrian hamsters (4-6 weeks old; ENVIGO) were intranasally inoculated with either the HPAIV H5N1 Bovine, Mountain Lion, or Human isolate. I.N. inoculation was performed with 10^4^ TCID_50_ of each isolate in 40 µL sterile DMEM. Post-inoculation, all animals were weighed daily and monitored for clinical signs of disease. Oropharyngeal and rectal swabs were collected daily to assess the virus shedding dynamics. At 10^4^ TCID50 of each H5N1 isolate in 40 µL sterile DMEM, six hamsters from each group were euthanized and respiratory tract tissues were collected. Six remaining animals were monitored for survival until 21 days and blood samples were collected after euthanasia.

To address the histopathological changes associated with Bovine, Mountain Lion, or Human isolate, four male and female Syrian hamsters (4-6 weeks old; ENVIGO) were inoculated I.N with 10^4^ TCID_50_ of each H5N1 isolates in 40 µL sterile DMEM. 4 dpi, respiratory tissues including NT, trachea and lungs were harvested. H&E staining and IHC analysis were performed to understand the histopathological changes and the virus distribution.

### Transmission study

Specially designed cages were used to perform transmission studies. Transmission cages were divided by 3D-printed perforated plastic dividers to allow airflow from the inoculated to the naïve hamster but prevent direct contact and fomite transmission (*36*). Donor hamsters (n□=□8) were inoculated I.N. as described above with 10^4^ TCID_50_ of each HPAIV H5N1 isolate. On day 1 post inoculation, donor animals were placed on one side of the cage together with the naïve contact sentinels (n=8). Likewise, one naïve animal was placed on the other side of the cage and served as the airborne sentinel (n=8). The divider placed between the cage allowed continuous airflow from the infected animal to the naïve airborne sentinel. Hamsters were cohoused for 48□h. The following day (D3), donor animals were euthanized, and sentinel animals were rehoused into regular rodent cages. Oropharyngeal swabs were collected for eight days. On day 21, animals were euthanized, and blood samples were collected.

### Viral RNA detection by RT-qPCR

Viral RNA was detected by qRT-PCR. RNA was extracted from swabs using a QiaAmp Viral RNA kit (Qiagen) according to the manufacturer’s instructions. Lung and NT tissues were homogenized, and RNA was extracted using the RNeasy kit (Qiagen). Viral M gene-specific primers, Forward primer: AAGACCAATCCTGTCACCTCTGA, Reverse primer: CAAAGCGTCTACGCTGCAGTCC and Probe: FAM-TTTGTGTTCACGCTCACCGTGCC-TAMRA were used for the detection of viral RNA. RNA (5 μl) was tested using the TaqMan Fast Virus One-Step Master Mix (Applied Biosystems) and the QuantStudio 3 Flex Real-Time PCR System (Applied Biosystems) according to the manufacturer’s instructions. Dilutions of Influenza standards with known genome copies were run in parallel to prepare a standard curve and calculate copy numbers/mL or copy numbers/g. The detection limit for the assay was 5 copies/reaction, and samples below this limit were considered negative.

### Virus titration

Infectious virus present in the swab and tissue samples was quantified by endpoint titration of 10-fold dilutions on MDCK cells in 96-well plates. Tissue samples were homogenized in 1 mL media using TissueLyser II (Qiagen). Cells were washed twice prior addition of 10-fold serially diluted swab samples and incubated the cells for 3 days at 37 °C and 5% CO_2_. Three days post-incubation, the presence or absence of infectious virus was measured by a standard HA assay using turkey red blood cells (Innovative Research). RBCs were washed at least three times with PBS and diluted to 0.33% in PBS directly before use. 75 µL of 0.33% RBCs was added to 25 µL of virus and incubated for 1 hour at 4°C. Subsequently, wells were marked as either agglutinated or negative. Titers were calculated using the Spearman-Karber Method.

### Enzyme-linked immunosorbent assay

Nunc MaxiSorp flat bottom 96-well plates (ThermoFisher Scientific) were coated with 50 ng in 50 µl/well of influenza A H5N1 A/Vietnam/1203/2004 hemagglutinin (HA) protein (IBT Bioservices) and incubated overnight at 4°C. Next day, the plates were blocked with 100 µL of casein in PBS (ThermoFisher Scientific) for 1 hour and incubated with serially diluted hamster sera (1:100-1:512000, in duplicate) for 1.5 h at room temperature. Immunoglobulin G (IgG) antibodies were detected by using affinity-purified polyclonal antibody peroxidase-labelled anti-hamster IgG (Seracare CAT# 5220-0371) at a dilution of 1:2500 in casein followed by 3,3′,5,5′-Tetramethylbenzidine 2-component peroxidase substrate (Seracare, 5120–0047) and stop solution (Seracare, 5150-0021). The optical density at 450 nm (OD450) was measured. Serological analysis of samples from the transmission study was performed at a 1:100 dilution. The threshold for positivity was determined as the average plus three times the standard deviation of the negative control hamster sera.

### Virus neutralizing antibody assay

Irradiated serum samples were heat-inactivated, two-fold serially diluted, and incubated with 100 TCID□□ of corresponding influenza viruses for 1 hour at 37°C. The virus-serum mixtures were added to confluent MDCK cells in 96-well plates and incubated at 37°C and 5% CO□. Three days post-incubation, the presence or absence of infectious virus was measured by a standard HA assay using fresh turkey red blood cells (Innovative Research). Neutralization titers were defined as the reciprocal of the highest serum dilution at which hemagglutination was absent.

### Histology and Immunohistochemistry

Tissues were fixed in 10% neutral buffered formalin x2 changes for a minimum of 7 days. Tissues were placed in cassettes and processed with a Sakura VIP-6 Tissue Tek on a 12-hour automated schedule, with a graded series of ethanol, xylene, and PureAffin. Embedded tissues were sectioned at 5um and dried overnight at 42 °C before staining. Immunoreactivity was detected using Millipore Sigma Anti-Influenza A nucleoprotein antibody at a 1:12,000 dilution. Roche Tissue Diagnostics DISCOVERY Omnimap anti-rabbit HRP was used as a secondary antibody. For negative controls, replicate sections from each control block were stained in parallel following an identical protocol, with the primary antibody replaced by Vector Laboratories rabbit IgG at a 1:2500 dilution. The tissues were stained using the DISCOVERY ULTRA automated stainer (Ventana Medical Systems) with a Roche Tissue Diagnostics DISCOVERY purple kit.

### Statistical analysis

Statistics were performed and significance was calculated as indicated where appropriate using GraphPad Prism 10 Software. p-values less than 0.05 were considered significant.

## Acknowledgements

We thank Dr. Todd Davis at the Centers for Disease Control and Prevention (CDC), Decatur, Georgia, USA, for providing the Human HPAIV H5N1 isolate A/Texas/37/2024 (EPI_ISL_19027114). We also thank Dr. Richard Webby at St. Jude Children’s Research Hospital, Memphis, Tennessee, USA, and Dr. Andrew Bowman at The Ohio State University, Columbus, Ohio, USA, for providing the Bovine HPAIV H5N1 isolate A/bovine/Ohio/B24OSU-342/2024 (EPI_ISL_19178076). We thank Erika Schwarz, Aracely Ospina Lopez, Nathaniel Antonioli, and Enrico Di Castro Young of the Molecular Diagnostics Section at the Montana Veterinary Diagnostic Laboratory, as well as Jennifer Ramsay and Matthew Becker of the Montana Fish, Wildlife & Parks Wildlife Health Laboratory, for providing the mountain lion specimen. This enabled tissue collection and isolation of the Mountain Lion HPAIV H5N1 isolate A/mountain lion/Montana/1/2024 (EPI_ISL_19083124). We acknowledge Dr. Joydeep Nag, Florida Research and Innovation Centre, Cleveland Clinic Lerner Research Institute for providing critical comments and valuable suggestions during the preparation of this manuscript. We are grateful to the Office of the Chief, RML, NIAID, NIH for their support within the high containment facility; the animal care staff of the Rocky Mountain Veterinary Branch, NIAID, NIH for their assistance during the study; and the Visual and Medical Arts Section, RML, NIAID, NIH for their help in preparing the graphical illustration.

## Author contributions

Conceptualization: RKM, VJM

Methodology: RKM, KCY, VJM

Investigation: RKM, FK, JES, SG, JPS, AW, KC, BJS, CC, CS, GS, EdW, NvD, KCY, VJM

Visualization: RKM, GS, VJM

Supervision: VJM

Writing - original draft: RKM, GS, VJM

Writing - review & editing: all authors.

## Data and materials availability

All data supporting the findings of this study have been deposited on Figshare and will be made publicly available upon acceptance. A DOI will be provided in the final published version.

## List of supplementary materials

1. Table 1. Summary of the amino acid variation among the three isolates

2. Table 2. Histology and immunohistochemistry analysis

