## supplement files for "Increased contact transmission of contemporary Human H5N1 compared to Bovine and Mountain Lion H5N1 in a hamster model"

Table 1. Summary of the amino acid variation among the three isolates

| | PB2 | | | | | | | | | | | | | | PB1 | | | |
|---|---|---|---|---|---|---|---|---|---|---|---|---|---|---|---|---|---|---|
| | 58 | 353 | 362 | 441 | 489 | 607 | 615 | 627 | 631 | 649 | 663 | 667 | 676 | 682 | 392 | 464 | 667 | 697 |
| A/bovine/Ohio/B24OSU-342/2024,Genotype B3.13 ( EPI_ISL_19178076) | A | K | G | N | S | L | I | E | L | I | K | V | A | G | V | D | I | E |
| A/mountain lion/Montana/01/2024, Genotype B3.6 (EPI_ISL_19083124) | T | R | E | D | P | I | M | E | M | V | R | I | T | S | I | D | T | G |
| A/Texas/37/2024, Genotype B3.13 (EPI_ISL_19027114) | A | K | E | N | S | L | I | K | M | I | K | V | A | G | I | N | I | E |

| | PA | | | | | | | | | | | NP | | | | NA | | NS | | |
|---|---|---|---|---|---|---|---|---|---|---|---|---|---|---|---|---|---|---|---|---|
| | 58 | 85 | 142 | 277 | 343 | 350 | 352 | 441 | 558 | 608 | 665 | 52 | 105 | 230 | 433 | 382 | 389 | 7 | 40 | 85 |
| A/bovine/Ohio/B24OSU-342/2024,Genotype B3.13 ( EPI_ISL_19178076) | S | T | K | S | S | S | D | M | S | T | L | H | M | F | T | E | V | L | Q | P |
| A/mountain lion/Montana/01/2024, Genotype B3.6 (EPI_ISL_19083124) | G | A | K | P | A | N | E | V | L | S | L | Y | V | L | N | D | M | S | Q | S |
| A/Texas/37/2024, Genotype B3.13 (EPI_ISL_19027114) | G | A | E | S | S | S | D | M | S | T | M | H | M | F | T | E | V | L | R | P |

| | PB2-S1 | | | | | | | | PA-X | | | NS-1 |
|---|---|---|---|---|---|---|---|---|---|---|---|---|
| | 58 | 353 | 362 | 441 | 495 | 499 | 500 | 503 | 58 | 85 | 142 | 88 |
| A/bovine/Ohio/B24OSU-342/2024,Genotype B3.13 ( EPI_ISL_19178076) | A | K | G | N | I | R | S | D | S | T | K | T |
| A/mountain lion/Montana/01/2024, Genotype B3.6 (EPI_ISL_19083124) | T | R | E | D | V | Q | N | G | G | A | K | I |
| A/Texas/37/2024, Genotype B3.13 (EPI_ISL_19027114) | A | K | E | N | I | R | N | E | G | A | E | I |

Table 2. Histology and immunohistochemistry analysis of the individual animals (Hamster 1 – 12).

| | A/bovine/Ohio/B24OSU-342/2024 | | | | A/mountain lion/Montana/01/2024 | | | | A/Texas/37/2024 | | | |
|---|---|---|---|---|---|---|---|---|---|---|---|---|
| **Histopathology** | **H1** | **H2** | **H3** | **H4** | **H5** | **H6** | **H7** | **H8** | **H9** | **H10** | **H11** | **H12** |
| **Trachea** | | | | | | | | | | | | |
| Epithelial degeneration and necrosis | 3 | 3 | 3 | 1 | 0 | 0 | 1 | 3 | 2 | 3 | 4 | 1 |
| Neutrophilic tracheitis | 4 | 2 | 4 | 2 | 0 | 0 | 2 | 3 | 2 | 3 | 4 | 1 |
| **Lungs** | | | | | | | | | | | | |
| Necrosis and neutrophilic bronchiolitis | 4 | 0 | 2 | 2 | 0 | 0 | 0 | 0 | 2 | 0 | 2 | 0 |
| Interstitial pneumonia | 2 | 0 | 1 | 0 | 0 | 0 | 0 | 0 | 0 | 0 | 1 | 1 |
| Pulmonary edema | 0 | 0 | 0 | 0 | 0 | 0 | 0 | 0 | 0 | 0 | 0 | 0 |
| Alveolar leukocytes | 2 | 0 | 2 | 1 | 0 | 0 | 0 | 0 | 1 | 0 | 0 | 0 |
| **Turbinates** | | | | | | | | | | | | |
| Respiratory epithelium degeneration and necrosis | 0 | 0 | 0 | 0 | 0 | 0 | 0 | 0 | 1 | 1 | 0 | 0 |
| Olfactory epithelium degeneration and necrosis | 0 | 0 | 0 | 0 | 0 | 0 | 0 | 0 | 0 | 0 | 0 | 0 |
| Infiltrates neutrophilic | 0 | 0 | 0 | 0 | 0 | 1 | 0 | 0 | 1 | 2 | 0 | 0 |
| **Immunohistochemistry** | **H1** | **H2** | **H3** | **H4** | **H5** | **H6** | **H7** | **H8** | **H9** | **H10** | **H11** | **H12** |
| **Trachea** | | | | | | | | | | | | |
| Epithelium | 2 | 2 | 2 | 2 | 0 | 0 | 0 | 3 | 1 | 2 | 4 | 0 |
| Macrophages | 1 | 1 | 1 | 0 | 0 | 0 | 0 | 1 | 0 | 0 | 1 | 0 |
| **Cumulative score** | 3 | 3 | 3 | 2 | 0 | 0 | 0 | 4 | 1 | 2 | 5 | 0 |
| **Lungs** | | | | | | | | | | | | |
| Alveolar macrophages | 2 | 3 | 1 | 1 | 0 | 0 | 1 | 3 | 3 | 0 | 2 | 0 |
| Bronchiolar epithelium | 3 | 3 | 2 | 2 | 0 | 0 | 0 | 1 | 4 | 0 | 4 | 0 |
| Pneumocytes | 2 | 0 | 1 | 1 | 0 | 0 | 0 | 0 | 0 | 0 | 1 | 0 |
| **Cumulative score** | 7 | 6 | 4 | 4 | 0 | 0 | 1 | 4 | 7 | 0 | 7 | 0 |
| **Nasal turbinates** | | | | | | | | | | | | |
| Respiratory epithelium | 0 | 1 | 0 | 0 | 0 | 3 | 0 | 1 | 4 | 3 | 0 | 2 |
| Olfactory epithelium | 0 | 0 | 0 | 0 | 0 | 2 | 0 | 0 | 1 | 0 | 0 | 3 |
| Macrophages | 1 | 1 | 0 | 0 | 0 | 1 | 1 | 0 | 1 | 1 | 0 | 1 |
| **Cumulative score** | 1 | 2 | 0 | 0 | 0 | 6 | 1 | 1 | 6 | 4 | 0 | 6 |

0 = No lesions
1 = Minimal (1-10%)
2 = Mild (11-25%)
3 = Moderate (26-50%)
4 = Marked (51-75%)
5 = Severe (76-100%)

**IHC attachment**
0=none
1=rare/few
2=scattered
3=moderate
4=numerous
5=diffuse
